# Optimal control of immune checkpoint inhibitor therapy in a heart-tumour model

**DOI:** 10.1101/2024.09.16.613200

**Authors:** Solveig A. van der Vegt, Ruth E. Baker, Sarah L. Waters

## Abstract

Autoimmune myocarditis, or cardiac muscle inflammation, is a rare but frequently fatal side–effect of immune checkpoint inhibitors (ICIs), a class of cancer therapies. Despite the dangers that side-effects such as these pose to patients, they are rarely, if ever, included explicitly when mechanistic mathematical modelling of cancer therapy is used for optimization of treatment. In this paper, we develop a two-compartment mathematical model which incorporates the impact of ICIs on both the heart and the tumour. Such a model can be used to inform the conditions under which autoimmune myocarditis may develop as a consequence of treatment. We use this model in an optimal control framework to design optimized dosing schedules for three types of ICI therapy that balance the positive and negative effects of treatment. We show that including the negative side-effects of ICI treatment explicitly within the mathematical framework significantly impacts the predictions for the optimized dosing schedule, thus stressing the importance of a holistic approach to optimizing cancer therapy regimens.

## 1 Introduction

As cancer treatment improves and patient survival increases, the long-term effects of treatments such as immune checkpoint inhibitors (ICIs) become an increasing concern [20]. Approximately 86-96% of patients treated with ICIs experience autoimmune side-effects, some of which can have life-threatening consequences [19, 41]. One such side-effect is autoimmune myocarditis, which occurs in 0.04-1.4% of patients treated with ICIs and proves fatal in 25-50% of cases [7, 26, 47]. Yet, despite these concerning statistics, the side-effects of cancer therapy are, to the best of our knowledge, not considered explicitly when mathematical models of cancer therapy are used for optimization of treatment.

The immune response consists of two pathways: the fast and non-specific innate immune response, and the slowly developing but antigen-specific adaptive immune response [36]. The adaptive immune response in turn consists of two pathways: one driven by T cells, and one driven by B cells. The autoimmune reaction in myocarditis is considered to be primarily driven by CD4+ T cells, so we focus on these T cells in our discussion and modelling of the adaptive immune response behind this disease [4]. The T-cell response consists of many different types of T cells with different functions that may be primarily pro-inflammatory, such as T helper (Th) cells, or anti-inflammatory, such as T regulatory (Treg) cells. An appropriate balance between the activity of pro- and anti-inflammatory immune cells is crucial in insuring that the immune response does not cause excessive damage to healthy tissue [36]. Autoimmune myocarditis can be caused by an injury to cardiac muscle tissue, for example due to a viral infection or cross-reacting T cells, which damages cells and exposes self-antigens that in turn trigger an autoimmune response [8]. Antitumour T cells that cross-react with cardiac antigens are a known potential trigger for autoimmune myocarditis as a side-effect of ICIs [30].

ICIs such as nivolumab and ipilimumab are therapeutic antibodies that block interactions that suppress the immune response. Ipilimumab is an antibody against cytotoxic T-lymphocyte-associated protein 4 (CTLA4), a protein expressed constitutively in high concentrations on the surface of Treg cells [4]. CTLA4 binds the surface proteins CD80 and CD86 found on dendritic cells, and these binding events lead to inhibition of T cell activation and inhibition of Treg cell depletion. The presence of ipilimumab blocks CTLA4 from binding, thus downregulating these inhibitory processes and stimulating the immune response [44]. Nivolumab is an antibody against programmed cell death protein-1 (PD-1). Blocking PD-1 from binding its ligands with nivolumab leads to increased activation of T cells and inhibition of their deactivation [4, 34]. Applying ICIs thus stimulates the immune response, which helps treat the tumor but can also affect other organs, leading to side-effects such as autoimmune myocarditis. For more details on the biology behind autoimmune myocarditis and the mechanisms of action of nivolumab and ipilimumab, we refer the interested reader to our recent review paper [42].

We recently proposed the first mathematical model of autoimmune myocarditis and the effects of the ICIs nivolumab and ipilimumab on the development and progression of this disease [43]. We now extend this model by coupling it to a simple tumour growth model, thus allowing us to consider the positive and negative effects ICIs simultaneously. We use the extended model to answer the following question: in a scenario where autoimmune myocarditis occurs as a side-effect of standard-of-care ICI treatment, can we design ICI dosing schedules that prevent autoimmune myocarditis from developing while maintaining inhibition of tumour growth? We turn to optimal control theory to explore what such treatment schedules might look like.

Optimal control theory uses control variables to optimize the output of a dynamical system by maximizing or minimizing a payoff function [21]. The theory has been used to predict, for example, optimal regimens for treating leukemia, minimizing autophagy and eradicating gliobastoma cells [25, 31, 32, 35, 38, 39]. In these studies, the control variable is often chosen to be the concentration of a drug in the patient’s body, or the rate at which the drug is administered. The two most common types of control used in designing optimal treatments are continuous and bang-bang controls. Although these types of controls are relatively straightforward to design, they lack biological realism. In the former case, the control variable is continuous in time. If the control variable is the concentration of a drug in a patient’s body, then continuous control is not realistic in most clinical settings as the drug concentration in the body cannot be tightly controlled. Bang-bang controls allow the control variable to switch between two distinct values, usually zero and some positive number. If the control variable is taken to be the influx rate of drug into the patient, this set-up is closer to how treatment is administered in the clinic, with the patient either receiving the drug, or not receiving the drug, at each point in time. However, ICI treatments are generally administered over a period of 30 to 90 minutes, and because bang-bang controls allow solutions where treatments are “on” for long periods of time, this type of control can also fail to provide clinically relevant solutions [15, 16, 38].

To obtain realistic treatment schedules, we select an approach in which, instead of optimizing the drug concentration or influx rate over time, we optimize the time points at which a drug dose of a set magnitude is administered [9]. Whereas continuous and bang-bang control are widely used, this approach has, to our knowledge, only been used previously in a handful of biomedical contexts [9, 10, 31].

In Section 2, we introduce the two-compartment mathematical model consisting of a heart compartment, captured via an adapted version of our previously developed model of autoimmune myocarditis, and a tumour compartment, defined by a simple tumour growth model. We explore the steady states of this model, and the behaviour of the two-compartment system under standard-of-care ICI treatment. Both nivolumab and ipilimumab can be administered individually as monotherapy, or together in combination therapy. We consider all three options: nivolumab monotherapy, ipilimumab monotherapy and combination therapy. In Section 3, we outline the optimal control algorithm used to optimize ICI dosing schedules. In Section 4, we use the optimal control framework in combination with the two-compartment model to optimize the treatment schedules of nivolumab monotherapy, ipilimumab monotherapy and combination therapy. Lastly, in Section 5, we summarize the results and discuss potential future avenues of research.

## 2 Two-compartment heart-tumour model

We now introduce the two-compartment model which describes the effects of the ICIs nivolumab and ipilimumab on the heart as well as on the tumour. The first compartment, representing the heart, consists of an adapted version of the model of autoimmune myocarditis we previously proposed [43]. The second compartment, representing the tumour, consists of a simple model of tumour growth. The primary aim of this work is to construct a framework for the optimization of ICI treatment schedules that consider both the positive and negative effects of ICIs. To this end, we present a simple model of tumour growth here which may be replaced by more complex and biologically realistic models in future work.

### 2.1 A model of autoimmune myocarditis

In a recent paper, we introduced the first mathematical model of autoimmune myocarditis and the effects of ICIs on the immunological mechanisms underlying this disease [43]. The well-mixed model describes the dynamics of damaged/dead cardiomyocytes (*C*), innate immune cells (*I*), pathogenic CD4+ T cells (*P*) and regulatory T cells (*R*) over time (*t*). Here, we extend this model by adding two ODEs describing the evolution of the mass (in grams) of nivolumab (*D*_*N*_) and ipilimumab (*D*_*I*_) in the body over time, as well as explicitly describing how the ICIs affect the rate at which some immunological process occur. The governing equations are

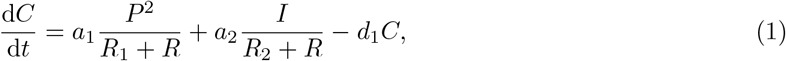

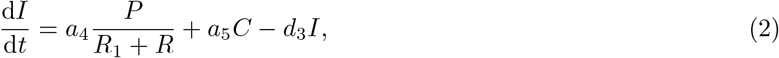

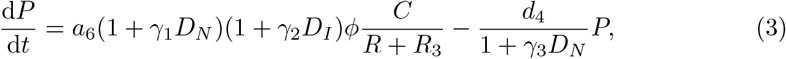

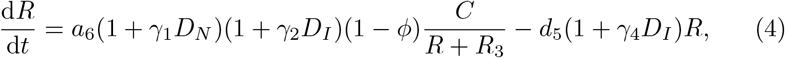

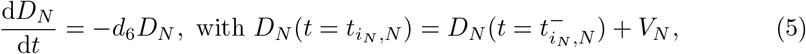

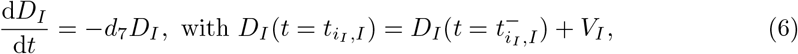

where all parameters are positive. The system is closed by specifying the initial conditions [*C*(0), *I*(0), *P* (0), *R*(0), *D*_*N*_ (0), *D*_*I*_ (0)] = [*C*_0_, *I*_0_, *P*_0_, *R*_0_, *D*_*N*,0_, *D*_*I*,0_].

In this model, the rates of T cell activation, pathogenic CD4+ T cell deactivation, and regulatory T cell deactivation are affected by the presence of ICIs. The parameters *a*_6_, *d*_4_ and *d*_5_ are the positive, ICI-free rates of T cell activation, pathogenic CD4+ T cell deactivation, and regulatory T cell deactivation, respectively, as defined in [43]. We define *V*_*N*_ and *V*_*I*_ as the fixed dose sizes of nivolumab and ipilimumab, respectively, both in grams. The changes in *D*_*N*_ and *D*_*I*_ when a dose of ICI is administered are assumed to be instantaneous as the time it takes to administer a dose of nivolumab or ipilimumab (30-90 minutes) is short compared to the length of the treatment window which is typically on the order of weeks or months [15, 16, 27]. We further define 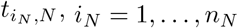, and 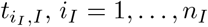, as the time points at which a dose of nivolumab or ipilimumab, respectively, are applied to system. The parameters *n*_*N*_ and *n*_*I*_ are fixed throughout the optimization. The parameters *d*_6_ *>* 0 and *d*_7_ *>* 0 are the decay rates of nivolumab and ipilimumab, respectively.

Both nivolumab and ipilimumab affect multiple immunological processes. However, to date the quantitative impact of each of the ICIs on the rates of T cell activation and inhibition are unknown. We therefore adopt a simplest-first approach and introduce the parameters *γ*_*i*_ ≥ 0, *i* = 1,…, 4, with units grams^−1^, which capture the pharmacodynamic impact of ICIs by translating the mass of ICIs in the body to fractional increases or decreases in the rate parameters associated with T activation and inhibition.

The values of *γ*_1_ and *γ*_3_ represent the impact of nivolumab on T cell activation and pathogenic CD4+ T cell deactivation, respectively, and the values of *γ*_2_ and *γ*_4_ represent the impact of ipilimumab on T cell activation and regulatory T cell deactivation, respectively. While the rarity of autoimmune myocarditis as a side-effect of ICIs makes estimates of incidence rates uncertain, the incidence rate of autoimmune myocarditis as a side-effect of combination therapy (0.27%) compared to monotherapy (0.06-0.09%) suggests a nonlinear synergistic effect on the immunological mechanisms underlying the disease when both ICIs are administered [2, 26]. To reflect this, we multiply the drug terms where these ICIs affect the same process, *i*.*e*. in the activation of T cells.

Our recent paper demonstrated that this model of autoimmune myocarditis has three steady states under ICI-free conditions: a “healthy” stable steady state at the origin which represents myocarditis-free conditions with zero activated immune cells and damaged cardiomyocytes present, a “diseased” stable steady state with high counts of immune cells and damaged cardiomyocytes which represents fully developed myocarditis, and a saddle point, which defines the threshold above which myocarditis develops. As ICI-therapy is administered, the saddle point moves closer to the origin, thus lowering the threshold for myocarditis to develop [43]. We consider that autoimmune myocarditis has developed if, at the final time point of the simulation, the cell counts in the system are above the saddle point as this guarantees that the system will converge to the “diseased” steady state.

### 2.2 A simple model of tumour growth

When it comes to mathematical modelling of tumour growth, the literature is vast, and many tumour growth laws have been proposed over the years [28, 45]. As different types of tumours follow different growth curves under different experimental conditions, no choice of growth law is universally applicable and thus a variety have been used in modelling cancer progression [24, 33, 37, 46].

The elimination of small tumours by the immune system, and the growth of those tumours which manage to escape it [5, 6, 40], suggests that tumour growth could be described by a mathematical model with three non-negative steady states: (1) a stable state at the origin which represents a healthy, cancer-free state; (2) a stable steady state which represents a tumour that has escaped elimination by the immune system and grown to carrying capacity; and (3) a saddle point which represents the upper bound on the initial tumour size that the immune system can then go on to eliminate. As nivolumab and ipilimumab stimulate the immune response against the tumour, this should enable the immune system to eliminate larger tumours, or at least to delay their growth. The presence of ICIs would thus be expected to lead to an increase in the tumour size at the saddle point. This is illustrated in Fig. 1.

**Fig. 1:**
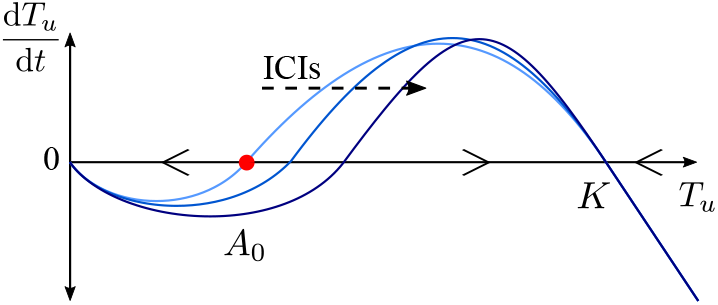
The RHS of Eq. (7) plotted as a function of the size of the tumour. For tumour volumes initially below the saddle point, *A*_0_ (red dot), the RHS of Eq. (7) is negative and tumour volume will decrease. For tumour volumes initially above the saddle point, the RHS of Eq. (7) is positive and the tumour will grow to carrying capacity, *K*. Increasing the level of ICIs in the system (dashed arrow) increases the tumour volume at the saddle point. For tumour volumes initially above the carrying capacity, the RHS of Eq. (7) is negative and tumour volume will decrease.

We denote tumour volume in cubic centimeters by *T*_*u*_, and propose a simple model of tumour dynamics of the form

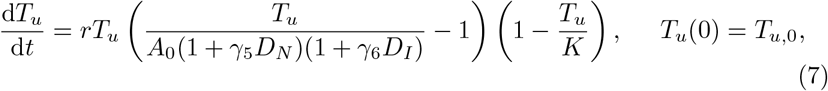

where the parameters *r, A*_0_, *γ*_5_, *γ*_6_, and *K* are positive constants with *r* denoting the growth rate of the tumour, *A*_0_ the threshold volume for elimination of the tumour by the immune system in an ICI-free environment, *K* the carrying capacity of the tumour, and *γ*_5_ and *γ*_6_ capturing the pharmacodynamics of nivolumab and ipilimumab, respectively, where the presence of ICIs increases the tumour volume at the saddle point. Eq. (7) admits three steady states, denoted *T*_*u*,*_, which capture the behaviour described above: *T*_*u*,*_ = 0 cm^3^, which is a stable steady state; *T*_*u*,*_ = *A*_0_(1 + *γ*_5_*D*_*N*_)(1 + *γ*_6_*D*_*I*_) cm^3^, which is a saddle point; and *T*_*u*,*_ = *K* cm^3^, which is a stable steady state.

In this work, we assume that at the start of the simulation the tumour has already escaped the immune system and thus *T*_*u*_(0) *> A*_0_, and that no treatment has yet been given. As ICIs are administered, two things can happen: (i) the saddle point *T*_*u*,*_ = *A*_0_(1+*γ*_5_*D*_*N*_)(1+*γ*_6_*D*_*I*_) increases but does not become larger than *T*_*u*_(*t*) so d*T*_*u*_/d*t* remains positive and the tumour continues to grow, though at a slower rate; or (ii) the saddle point *T*_*u*,*_ = *A*_0_(1+*γ*_5_*D*_*N*_)(1+*γ*_6_*D*_*I*_) becomes larger than *T*_*u*_(*t*), so d*T*_*u*_/d*t* becomes negative and the tumour shrinks, at least initially. In reality, treatment with nivolumab and/or ipilimumab delays tumour growth but in most cases does not lead to tumour elimination [1, 17, 22]. For the remainder of this work, we thus ensure that we are in a regime where *T*_*u*_(0) is sufficiently large that d*T*_*u*_/d*t* is decreased by the presence of ICIs, but remains positive.

### 2.3 Coupling the heart and tumour compartments

To construct a model that captures the positive effects of ICIs on tumour growth and the negative effects of ICIs on the heart, we now couple the heart and tumour compartments by adding a small, unidirectional flow of pathogenic T cells from the tumour to the heart. We assume this flow to be unidirectional as the activation of cross-reactive pathogenic T cells initially happens as part of the anti-tumour immune response, and these activated pathogenic T cells can then circulate through the body and end up in the heart. The number of active T cells coming from the heart to the tumour microenvironment is assumed to be negligible compared to the number of active T cells already in the tumour microenvironment.

The unidirectional flow of pathogenic CD4+ T cells is represented in the model by the addition of a term to Eq. (3) which is highlighted here in red:

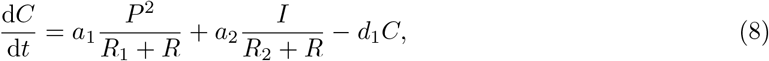

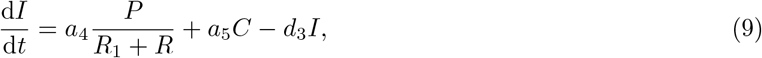

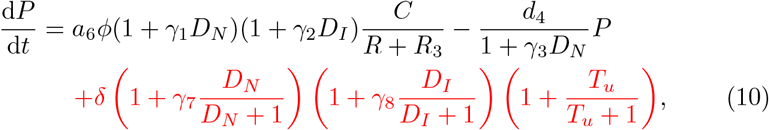

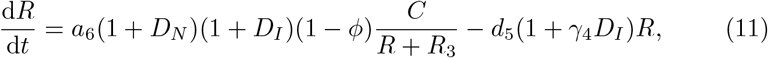

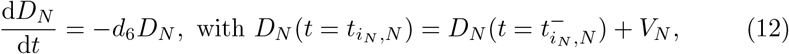

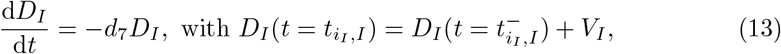

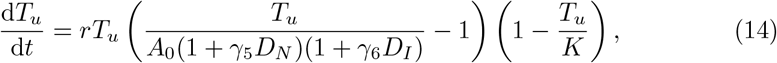

where *δ, γ*_7_ and *γ*_8_ are positive, constant parameters. For a schedule consisting of a fixed number of *n* = *n*_*N*_ + *n*_*I*_ total ICI doses, the time points 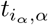, where *α* ∈ {*N, I*}, *i*_*N*_ = [1, …, *n*_*N*_] and *i*_*I*_ = [1, …, *n*_*I*_], are the time points at which a dose of ICIs is administered (nivolumab if *α* = *N* and ipilimumab if *α* = *I*). The parameter *δ* represents the baseline flow of cardiac-antigen specific T cells that circulate through the body in a healthy state, which is assumed to be small [18, 23]. The presence of a tumour is assumed to increase this flow as the tumour provides a source of activated cross-reacting T cells. The number of active T cells in the tumour compartment is assumed to correlate with tumour volume. The quantitative of effect of a large tumour on the flow activated cross-reactive T cells is currently unknown but assumed to be relatively small as the risk of autoimmune myocarditis for cancer patients not being treated with ICIs is small, hence we define a term which at most double the rate *δ*. Because nivolumab and ipilimumab both stimulate the anti-tumour immune response, the presence of these ICIs is assumed to further increase the level of T cells in the tumour compartment and thus in circulation throughout the body. The parameters *γ*_7_ and *γ*_8_ translate the masses of nivolumab and ipilimumab, respectively, in the system to the effect on the rate of T cell activation in the tumour compartment, which is assumed to be bound by a maximum rate at which T cells can be activated and enter circulation. The system of ODEs is closed by defining the initial conditions [*C*(0), *I*(0), *P* (0), *R*(0) *D*_*N*_ (0), *D*_*I*_ (0), *T*_*u*_(0)] = [*C*_0_, *I*_0_, *P*_0_, *R*_0_, *D*_*N*,0_, *D*_*I*,0_, *T*_*u*,0_].

### 2.4 Parameter values

It remains to specify parameter values for the two-compartment model. The parameter values previously defined for the mathematical model of autoimmune myocarditis are used again here and reproduced in Tbl. 2 [43]. In the heart compartment model, we need values for the decay rates of nivolumab, *d*_6_, and ipilimumab, *d*_7_, as well as the values of *γ*_*i*_, *i* = 1, …, 4. We set *d*_6_ = log(2)*/*25 days^−1^ as the half-life of nivolumab in the body is 25 days [11]. Similarly, we set *d*_7_ = log(2)*/*14.7 days^−1^ as the half-life of ipilimumab in the body is 14.7 days [11]. In the absence of specific values in the literature, we set *γ*_*i*_ = 1 grams^−1^ for *i* = 1, …, 4.

We must also specify values for the parameters in the tumour growth model, *r, A*_0_, *K, γ*_5_ and *γ*_6_, and the parameters which govern the flow of pathogenic T cells: *δ, γ*_7_ and *γ*_8_. In the absence of data, we assign for these parameter values such that the assumptions made in the previous sections are not violated, and such that the model shows the required qualitative behaviour as we would expect based on the literature. The parameters *r, A*_0_ and *K* govern the growth of the tumour, and *γ*_5_ and *γ*_6_ translate the masses of nivolumab and ipilimumab, respectively, to their effect on the tumour. In the absence of ICIs a tumour with a size greater than *A*_0_ should grow to carrying capacity whereas the presence of nivolumab or ipilimumab should delay the growth of the tumour by a period on the order of months to match with the average progression-free survival timescale of patients treated with these ICIs [1, 17, 22]. In addition, we must take *δ* to be sufficiently small that autoimmune myocarditis does not develop in an ICI-free environment, regardless of tumour volume, as there is no evidence that the presence of a tumour alone largely increases the risk of autoimmune myocarditis developing.

Patients do not recover from autoimmune myocarditis by simply stopping treatment with ICIs. This is reflected in our model by the fact that once autoimmune myocarditis has developed, *i*.*e*. the system is at or approaching the stable steady state with high levels of immune cells, the system cannot return to the healthy steady state even in an ICI-free environment. We assume here that an adjustment to the dosing schedule can be made at the earliest eight days after the first dose of ICIs has been administered, as the T cell immune response takes approximately seven days to fully develop and hence this is the period of time that must elapse before a pathogenic response to ICI therapy can be detected [36]. If the increase in inflammation after the first dose of ICIs is so large that autoimmune myocarditis immediately develops, it is not possible to avoid this side-effect by optimizing subsequent doses. Therefore, to be in a position where we can use optimal control theory to adjust dosing schedules to prevent autoimmune myocarditis, we must be in a scenario where myocarditis develops as a consequence of the second or later doses. We thus require the values of *γ*_7_ and *γ*_8_ to be such that when standard-of-care treatment is applied there are values of *V*_*N*_ and *V*_*I*_, the drug doses of nivolumab and ipilimumab, respectively, for which autoimmune myocarditis develops for nivolumab monotherapy, ipilimumab monotherapy as well as combination therapy, but at the earliest after the second dose.

**Table 1:** Standard-of-care dosing schedules for nivolumab and ipilimumab monotherapy, and combination therapy.

| type of therapy | Dosing days | Reference |
| --- | --- | --- |
| Nivolumab monotherapy | 2, 16, 30, 44, 58 | [15] |
| Ipilimumab monotherapy | 2, 23, 44, 65 | [16] |
| Combination therapy | 2, 23, 44, 65, 86 (nivolumab only) | [15, 16] |

Whether we are in such a scenario thus also depends on the values we specify for the drug doses: *V*_*N*_ for nivolumab and *V*_*I*_ for ipilimumab. Tbl. 1 shows the standard-of-care schedules for nivolumab and ipilimumab monotherapy, and combination therapy that we used in this work. The dose of nivolumab is the same for combination therapy and monotherapy, but the dose of ipilimumab in combination therapy is, on average, approximately one tenth of its value is in monotherapy [15, 16]. We assume that doses of ICIs are administered instantaneously at the time points specified by the dosing schedule.

For the parameter values specified in Tbl. 2, with *V*_*N*_ = 0.25 grams for nivolumab monotherapy and *V*_*I*_ = 1.0 grams for ipilimumab monotherapy, we see the desired behaviour when standard-of-care treatment is applied, where tumour growth is delayed but autoimmune myocarditis develops after the second dose or later (see Fig. 2). In these simulations, we assume that the heart is healthy and ICI-free initially and that the tumour has an initial volume of 0.1 cm^3^, so the initial conditions are [*C*_0_, *I*_0_, *P*_0_, *R*_0_, *D*_*N*,0_, *D*_*I*,0_, *T*_*u*,0_] = [0, 0, 0, 0, 0, 0, 0.1]. We choose this initial tumour volume to use throughout this work as it is well above the ICI-free saddle point of 0.001 cm^3^ and the tumour will thus grow initially even in scenarios where ICIs are applied. In Fig. 2, we show that in ICI-free conditions, the tumour grows to carrying capacity quickly and the level of damaged cardiomyocytes in the heart remains low, indicating that myocarditis does not develop. The simulations in the bottom three rows of Fig. 2 show that for all three types of ICI therapy, we see a delay in tumour growth, but autoimmune myocarditis develops as a side-effect of treatment, as indicated by the high levels of damaged cardiomyocytes in the system at the final time point. Comparing tumour growth in top row of Fig. 2 to tumour growth in the other three rows, we see that tumour growth is delayed on the order of months when ICIs are administered. This is consistent with observed patient responses to ICI therapy, so the simple tumour model as presented here shows the desired qualitative behaviour [1, 17, 22].

**Fig. 2:**
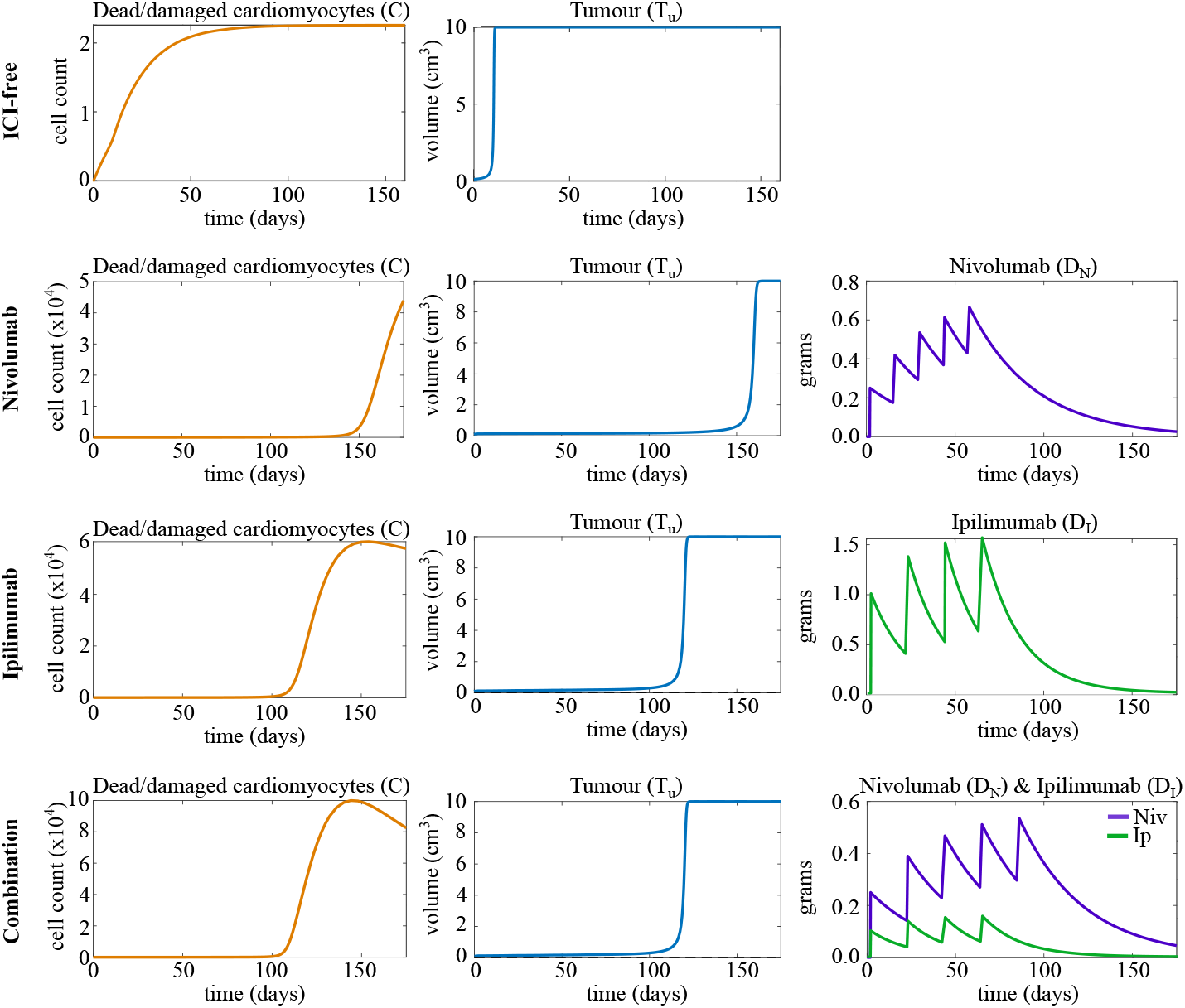
Time traces of some state variables under ICI-free conditions (top), with nivolumab, with ipilimumab and with combination therapy (bottom). Therapies are applied according to their respective standard-of-care schedules (Tbl. 1). The initial tumour volume is 0.1 cm^3^, and all other variables are initialized at zero. We set *V*_*N*_ = 0.25 grams and *V*_*I*_ = 0.1 grams, and all other parameter values are as listed in Tbl. 2.

Thus, with the parameter values established here we are in a parameter regime where we can use optimal control theory to investigate the problem of balancing tumour treatment with myocarditis prevention. We note, however, that the selected parameter set that gives the desires behaviour is non-unique, and other combinations of parameter values are possible. As quantitative experimental data becomes more available the selection of parameter values can be refined.

## 3 Optimal control of timing of doses

We now set out the algorithm developed by Castiglione and Piccoli that we use to optimize the timing of the ICI doses to ensure that tumour growth is delayed while also ensuring that autoimmune myocarditis does not develop [9, 29]. This method relies on finding the optimal schedule by minimizing a payoff function through gradient descent, and applies to a system of ODEs of the form

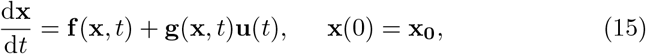

where **x**(*t*) are the state variables and **u**(*t*) are the control variables. For the purpose of optimizing ICI treatment schedules to prevent autoimmune myocarditis, we define the function **f** (**x**, *t*) as the right-hand side (RHS) of Eq. (8)-(14), the vector **x** = [*C, I, P, R, D*_*N*_, *D*_*I*_, *T*_*u*_], and the control

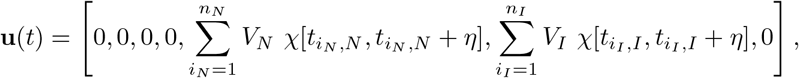

where *n*_*N*_ is the number of nivolumab doses in the schedule, *n*_*I*_ is the number of ipilimumab doses in the schedule, *χ* is an indicator function which is equal to one when 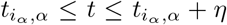 where *α* ∈ {*N, I*} and zero otherwise. The parameter *η* is the duration over which an ICI dose is administered, which is assumed to be negligibly small compared to the length of the treatment window [9]. Thus we will assume instantaneous dosing at the time points specified in the dosing schedule for the remainder of this work. Lastly, we set **g**(**x**, *t*) to be the identity matrix. The payoff function in the algorithm develop by Castiglione and Piccoli takes the form

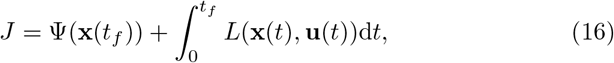

where *t*_*f*_ is the final time point of the simulation. This payoff function allows the user of the algorithm to consider both the state of the system at the final time point as well as the behaviour of the system over time in the computation of the payoff.

**Table 2:** Definitions, values and units of model parameters.

| Param. | Definition | Value | Units |
| --- | --- | --- | --- |
| $a_1$ | Rate at which pathogenic CD4+ T cells damage/kill cardiomyocytes | 80 | day<sup>-1</sup> |
| $a_2$ | Rate at which innate immune cells damage/kill cardiomyocytes | 6000 | cells day<sup>-1</sup> |
| $a_4$ | Rate at which pathogenic CD4+ T cells activate innate immune cells | 500,000 | cells day<sup>-1</sup> |
| $a_5$ | Rate at which damaged/dead cardiomyocytes activate innate immune cells | 35 | day<sup>-1</sup> |
| $a_6$ | Rate at which damaged/dead cardiomyocytes activate pathogenic CD4+ T cells | 750 | cells day<sup>-1</sup> |
| $d_1$ | Clearance rate of damaged/dead cardiomyocytes | 75 | day<sup>-1</sup> |
| $d_3$ | Innate immune cell death rate | 7 | day<sup>-1</sup> |
| $d_4$ | Pathogenic CD4+ T cell death rate | 0.1 | day<sup>-1</sup> |
| $d_5$ | Regulatory T cell death rate | 0.1 | day<sup>-1</sup> |
| $d_6$ | Degradation rate of nivolumab | log(2)/25 | day<sup>-1</sup> |
| $d_7$ | Degradation rate of ipilimumab | log(2)/14.7 | day<sup>-1</sup> |
| $R_1$ | Number of regulatory immune cells for 50% inhibition of pathogenic CD4+ T cell activity | 1100 | cells |
| $R_2$ | Number of regulatory immune cells for 50% inhibition of innate immune cells | 40000 | cells |
| $R_3$ | Number of regulatory immune cells for 50% inhibition of pathogenic CD4+ T cell activation | 2000 | cells |
| $\phi$ | Fraction activated CD4+ T cells that take on a pathogenic phenotype | 0.7 | unitless |
| $\delta$ | Rate of transport of T cells from tumour to heart | 0.5 | cells day<sup>-1</sup> |
| $r$ | Maximal growth rate of the tumour | 0.001 | cm<sup>3</sup> day<sup>-1</sup> |
| $A_0$ | Maximum tumour volume for immune system control | 0.001 | cm<sup>3</sup> |
| $K$ | Carrying capacity of tumour | 10 | cm<sup>3</sup> |
| $\gamma_1$ | Relative effect of nivolumab on T cell activation | 1 | grams<sup>-1</sup> |
| $\gamma_2$ | Relative effect of ipilimumab on T cell activation | 1 | grams<sup>-1</sup> |
| $\gamma_3$ | Relative effect of nivolumab on pathogenic T cell death | 1 | grams<sup>-1</sup> |
| $\gamma_4$ | Relative effect of ipilimumab on regulatory T cell death | 1 | grams<sup>-1</sup> |
| $\gamma_5$ | Relative effect of nivolumab on T cell activation and transport from tumour site | 100 | grams<sup>-1</sup> |
| $\gamma_6$ | Relative effect of ipilimumab on T cell activation and transport from tumour site | 25 | grams<sup>-1</sup> |
| $\gamma_7$ | Relative effect of nivolumab on T cell activation against the tumour | 5 | grams<sup>-1</sup> |
| $\gamma_8$ | Relative effect of ipilimumab on T cell activation against the tumour | 5 | grams<sup>-1</sup> |

Below we set out the algorithm for optimizing dosing schedules that consist of *n*_*N*_ doses of nivolumab and *n*_*I*_ doses of ipilimumab. The dosing schedule 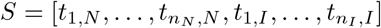 specifies the time points at which doses of ICIs are administered. The total number of doses *n*_*N*_ +*n*_*I*_ is fixed, but multiple doses can be administered at the same time point. A time point may thus appear in *S* multiple times, indicating a larger dose being administered at that time point. We define *S*_*j*_ to be the dosing schedule in *j*-th iteration, where *j* = 0, … and *S*_0_ corresponds to the initial dosing schedule. The algorithm proceeds as follows:

1. Set the ICI dose sizes *V*_*N*_ and *V*_*I*_, the initial dosing schedule 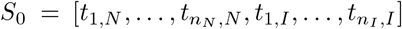 for *n* = *n*_*N*_ + *n*_*I*_ total doses to be administered, the length of the treatment window [*t*_min_, *t*_max_], the final time point of the simulation *t*_*f*_, the gradient descent step size parameter *h*, the initial conditions **x**_**0**_, and the convergence threshold for the payoff *c*_*p*_.
2. For the (*j* +1)-th iteration, solve the ODEs in Eq. (8)-(14) using the current schedule *S*_*j*_ to obtain the time traces **x**(*t*) of the state variables.
3. For each 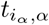 in *S*_*j*_ where *α* ∈ {*N, I*}, *i*_*N*_ ∈ {1, …, *n*_*N*_ }, and *i*_*I*_ ∈ {1, …, *n*_*I*_ }, solve the following variation equations over the time span 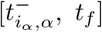

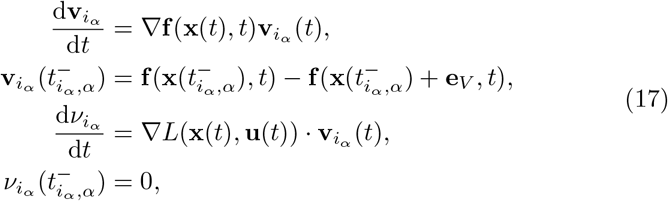

where the matrix ∇**f** and the vector ∇*L* denote the Jacobians of **f** and *L*, respectively, and 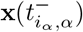 denotes the state of the system immediately before the dose is given. The vector **e**_*V*_ denotes the ICI dose that is given at time 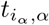, *i*.*e*. **e**_*V*_ = [0, 0, 0, 0, *V*_*N*_, 0, 0] if *α* = *N*, and **e**_*V*_ = [0, 0, 0, 0, 0, *V*_*I*_, 0] if *α* = *I*.
4. For each 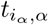 in *S*_*j*_, where *α* ∈ {*N, I*}, *i*_*N*_ ∈ {1, …, *n*_*N*_ }, and *i*_*I*_ ∈ {1, …, *n*_*I*_ }, compute the direction of steepest descent as

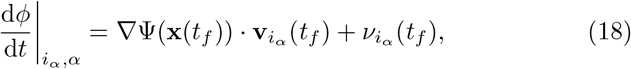

where the vector ∇Ψ denotes the Jacobian of Ψ.
5. Update the treatment schedule *S*_*j*_ as

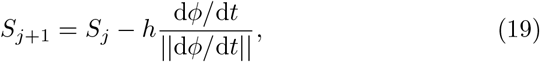

where

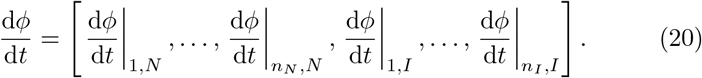

If any 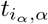 in *S*_*j*+1_ is before the first day of the treatment window, *t*_min_, or after the last day, *t*_max_, we set that 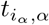 equal to *t*_min_ or *t*_max_, respectively, to ensure that all doses are applied within the allowed treatment window.
6. If *j* = 0, *i*.*e*. this is the first iteration, return to step 2. For *j >* 0, compute the payoff associated with the current schedule *S*_*j*_. If 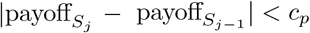, terminate the algorithm. If not, return to Step 2.

In Step 5, d*ϕ/*d*t* is divided by its norm to improve convergence: the values in the vector d*ϕ/*d*t* can vary widely, so dividing by the norm controls the size of the steps taken in the gradient descent algorithm. Other options for optimizing the gradient descent algorithm in Step 5 can be found in [12], however we found the approach in Eq. (19) sufficient from a computational perspective.

## 4 Optimization of ICI therapy

We now use the optimal control algorithm detailed in Section 3 to find a schedule of ICI doses that balances the need to prevent autoimmune myocarditis with the need to delay tumour growth. Our primary aim is to identify a schedule that prevents autoimmune myocarditis and delays tumour growth compared to no treatment. An optimized, though not necessarily globally optimal, schedule would thus suffice to achieve our objective and it is this we seek here.

To account for both the positive and negative effects of ICI treatment. We consider the payoff function of the form

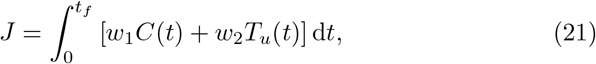

where *w*_*i*_ *>* 0 for *i* = 1, 2 are the weights that determine the relative importance of the terms in the payoff function. This payoff function ensures that a penalty is incurred if autoimmune myocarditis develops or the tumour grows quickly. An optimized dosing schedule will strike a careful balance between delaying tumour growth and preventing autoimmune myocarditis.

In each case, we initialize the model with an ICI-free, inflammation-free heart compartment and a growing tumour which is small compared to the carrying capacity, *i*.*e*. [*C*_0_, *I*_0_, *P*_0_, *R*_0_, *D*_*N*,0_, *D*_*I*,0_, *T*_*u*,0_] = [0, 0, 0, 0, 0, 0, 0.1]. We set the weights as *w*_1_ = 1 cells^−1^ and *w*_2_ = 1 cm^−3^ for simplicity. The convergence threshold is set to *c*_*p*_ = 0.1, and the gradient descent step size, *h*, is equal to 0.01. We set 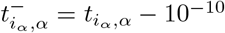.

The first dose of ICI therapy is administered on day 2, which is not part of the optimization. We let the treatment window start on day 10, so *t*_min_ = 10 days to reflect the time it takes for the immune response to develop and become clinically detectable. The upper bound on the treatment window varies by type of ICI therapy. The simulations are stopped 100 days after the end of the treatment window to ensure we capture the development of autoimmune myocarditis should it develop as a consequence of a dose on the last day of the treatment window. An overview of the timeline of the optimization problem is included in Fig. 3.

**Fig. 3:**
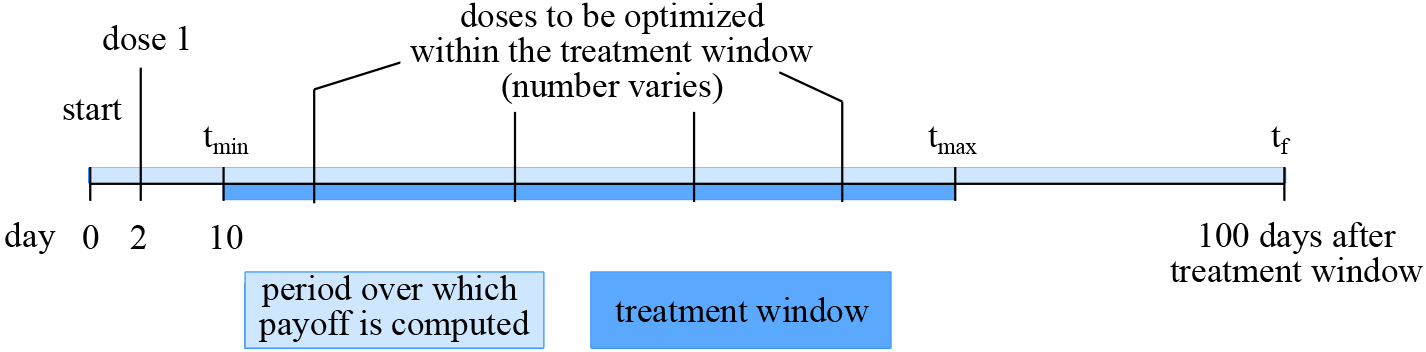
Timeline of the optimization problem for ICI treatment schedule design.

Our aim is to consider the effects of ICIs in the heart and tumour simultaneously. In the Supplementary Information, we isolate the tumour problem and optimize dosing schedules for tumour inhibition only. We find that dosing schedules optimized for tumour growth inhibition only do not differ significantly from the standard-of-care treatment (see Section 1 of the Supplementary Information). In Sections 4.1-4.3, we explore in turn the optimized treatment schedules for nivolumab monotherapy, ipilimumab monotherapy and combination therapy in the two-compartment model.

### 4.1 Nivolumab monotherapy

We initialize the dosing schedule at the standard-of-care schedule of nivolumab monotherapy, which is a dose every two weeks starting from day 2, so that *S*_0_ = [16, 30, 44, 58]. The final day of the treatment window is day 60 (*i*.*e. t*_max_ = 60 days), and *V*_*N*_ = 0.25 grams. The top two panels in Fig. 4 show the results of the optimization over the iterations of the gradient descent algorithm. The optimized dosing schedule is *S* = [10.0, 22.7, 59.7, 60.0].

**Fig. 4:**
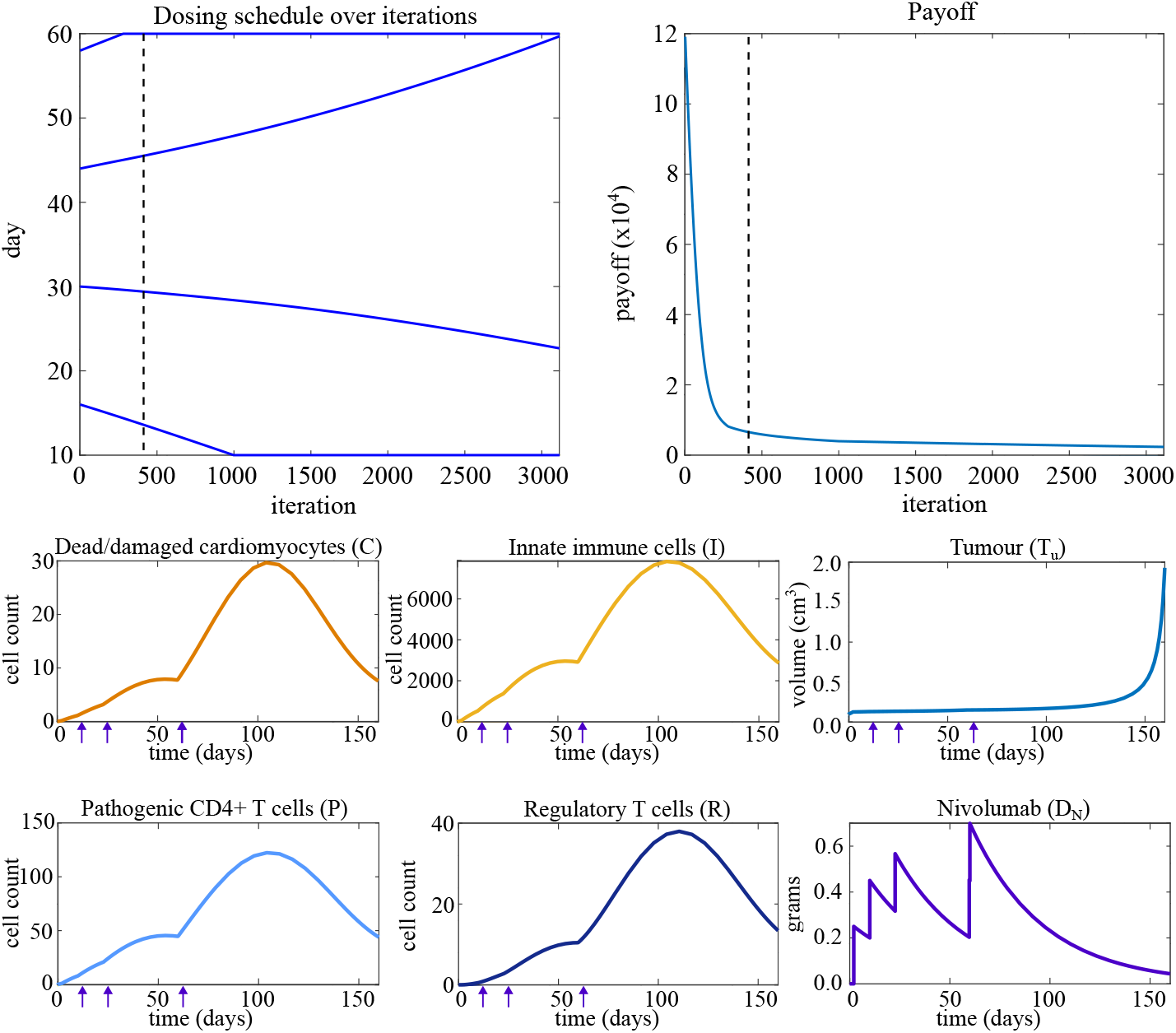
Top left: the dosing schedule of nivolumab monotherapy evolving over iterations of the optimization algorithm. Top right: the payoff evolving over iterations of the optimization algorithm. The dashed line indicates the first iteration at which autoimmune myocarditis does not develop. Bottom: Time traces of model variables when the optimized dosing schedule for nivolumab monotherapy is applied. Arrows indicate time points at which a dose of nivolumab is administered (first, not-optimized dose on day 2 not included).

The top left-hand panel of Fig. 4 shows that optimizing for the prevention of autoimmune myocarditis as well as delay of tumour growth delays the third and fourth doses while bringing doses one and two forward such that the treatment uses the whole treatment window. Regular dosing keeps tumour growth under control, while the increased time between the second and third doses compared to the standard-of-care schedule allows immune cell counts to decrease, which prevents autoimmune myocarditis from developing, as indicated by the low counts of immune cells at the final time point.

The bottom six panels in Fig. 4 show the time traces of the model variables for the optimized dosing schedule. Comparing the bottom plots in Fig. 4 to the second row of plots in Fig. 2 shows that the optimized treatment schedule prevents autoimmune myocarditis from developing without diminishing the effect of treatment on the tumour.

### 4.2 Ipilimumab monotherapy

When consideren ipilimumab monotherapy, we initialize the schedule at the standard-of-care treatment schedule, *S*_0_ = [23, 44, 65]. The final day of the treatment window (*t*_max_) is day 70, and *V*_*I*_ = 1.0 grams.

The optimized ipilimumab treatment schedule is *S* = [10.0, 47.4, 70.0]. When we consider the results of the optimization in the top panels in Fig. 5, we see a similar trend to what we saw for nivolumab monotherapy. Compared to the standard-of-care dosing schedule, the optimized dosing schedule spreads out the ICI doses over the entire dosing window. As shown in the bottom panels in Fig. 5, this optimized dosing schedule delays tumour growth comparably to the standard-of-care schedule (see third row in Fig. 2), but does not cause autoimmune myocarditis to develop. The optimized schedule thus balances the positive and negative effects of ICI therapy effectively.

**Fig. 5:**
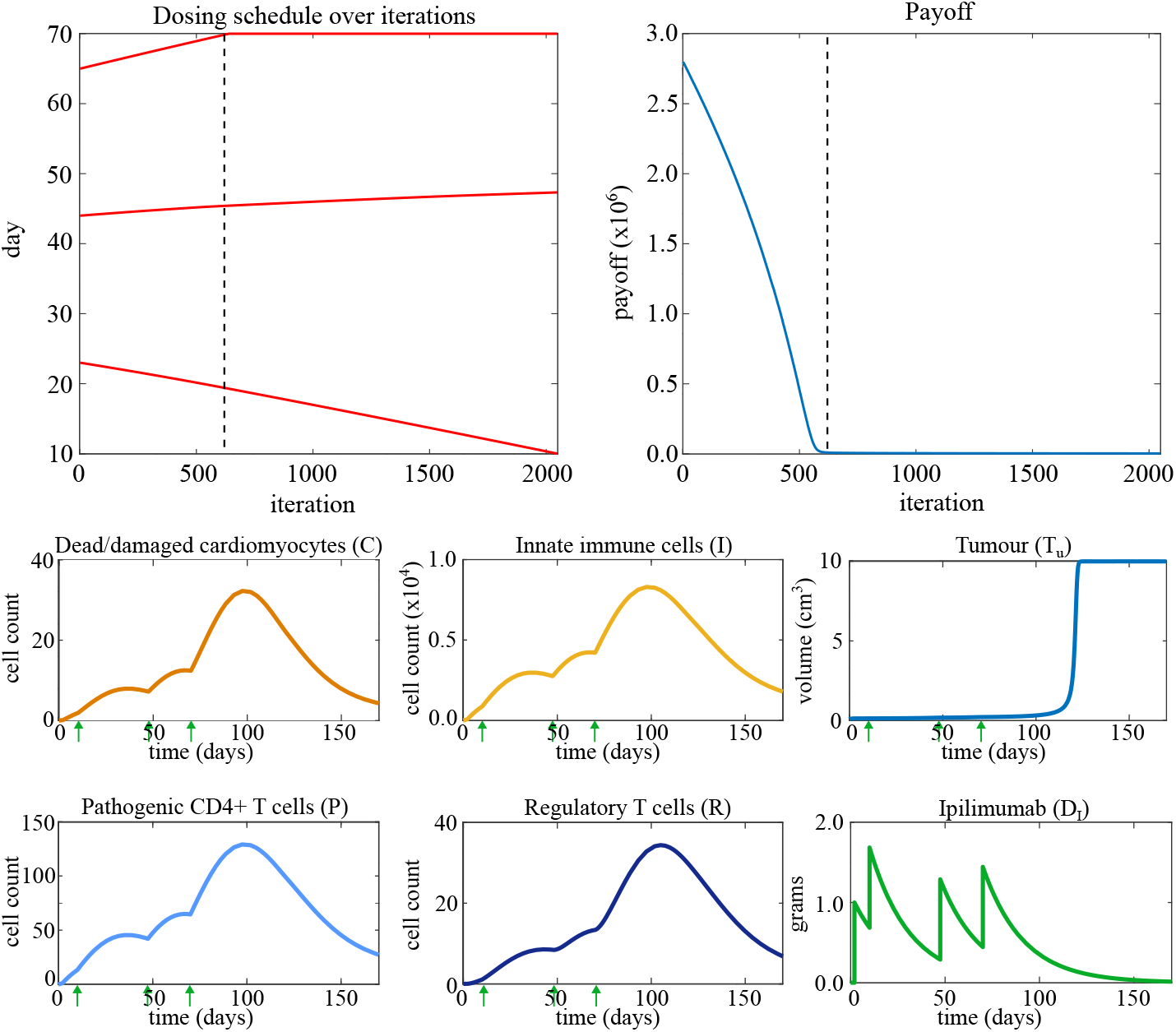
Top left: the dosing schedule of ipilimumab monotherapy evolving over iterations of the optimization algorithm. Top right: the payoff evolving over iterations of the optimization algorithm. The dashed line indicates the first iteration at which autoimmune myocarditis does not develop. Bottom: Time traces of model variables when the optimized dosing schedule for ipilimumab monotherapy is applied. Arrows indicate a time point at which a dose of nivolumab is administered (first, not-optimized dose on day 2 not included).

### 4.3 Combination therapy

Lastly, we turn to the optimization of treatment schedules for combination therapy. We use the standard-of-care schedule as a starting point for our optimization, *S*_*N*,0_ = [23, 44, 65, 86], and *S*_*I*,0_ = [23, 44, 65]. We set *V*_*N*_ = 0.25 grams and *V*_*I*_ = 0.1 grams, and the end of the treatment window (*t*_max_) at day 90. The gradient descent algorithm takes a long time to converge for this problem. To alleviate this issue we initially set *h* = 0.1 until the payoff drops below 10^4^ at which point we set *h* = 0.01.

The optimized dosing schedule is *S*_*N*_ = [10.0, 90.0, 90.0, 90.0] and *S*_*I*_ = [10.0, 24.9, 63.7]. The top panels in Fig. 6 shows the dosing schedule and the payoff over the iterations. Compared to the standard-of-care schedule shown in the bottom row of Fig. 2, the optimized schedule spreads out the ICI doses over the treatment window. In particular, we note a large amount of time between the dose of nivolumab on day 10 and the three doses of nivolumab administered simultaneously on day 90. The doses of ipilimumab are administered in between but, as these doses are relatively small in combination therapy, they do not cause large perturbations to the immune system. There is thus a lot of time in between the doses of nivolumab for the immune system to relax and the counts of immune cells in the heart to decrease. As shown in the bottom panels of Fig. 6, the optimized schedule does not cause autoimmune myocarditis to develop while maintaining inhibition of tumour growth on a similar scale to the standard-of-care.

**Fig. 6:**
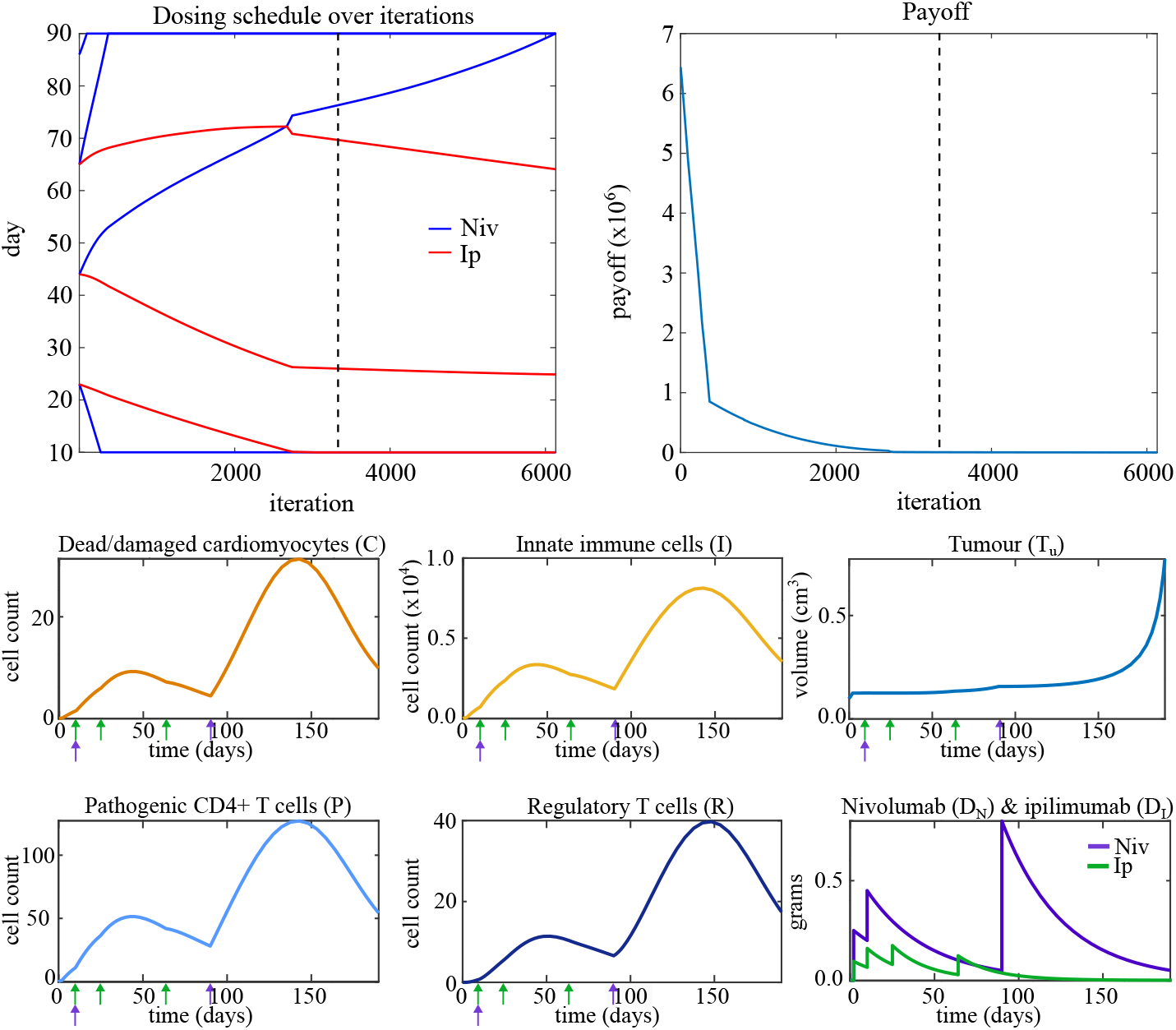
Top left: the dosing schedule of combination therapy evolving over iterations of the optimization algorithm. Top right: the payoff evolving over iterations of the optimization algorithm. The dashed line indicates the first iteration at which autoimmune myocarditis does not develop. Bottom: time traces of model variables when the optimized dosing schedule for combination therapy is applied. Arrows indicate time points at which a dose of nivolumab (purple) or ipilimumab (green) is administered (first, not-optimized doses on day 2 not included).

In summary, we have shown that consideration of the effects of ICIs on tumour growth in addition to the risk of autoimmune myocarditis significantly impacts predictions of safe and effective dosing schedules. When the only objective is to delay tumour growth, we recover the standard-of-care treatment schedule. When we also consider the prevention of autoimmune myocarditis, the ICI doses are better spread out over the treatment window, striking a careful balance between regular dosing, so as to control tumour growth, and keeping ICI levels low enough such that the stimulation of the immune system does not cause the development of autoimmune myocarditis.

In the Supplementary Information, we explore how the results discussed here change as the dose sizes *V*_*N*_ and *V*_*I*_, the length of the treatment window, the step parameter *h*, and the relative values of the payoff function weights *w*_1_ and *w*_2_ are varied for all three types of therapy. We show that, for all three types of therapy, there exists a (narrow) range of values of *V*_*N*_ and *V*_*I*_ for which autoimmune myocarditis develops under the standard-of-care dosing schedule, but can be prevented by an optimized schedule. We further show that the results are robust to changes in the length of the treatment window, the step size *h*, and the payoff function weights *w*_1_ and *w*_2_, although some combinations of values for *w*_1_ and *w*_2_ require careful calibration of the parameter *h* to ensure that the algorithm converges to a myocarditis-free solution when one exists.

## 5 Discussion

This work shows that in designing treatment regimens it is essential to consider the off-target effects of cancer treatment, as well as the impact of the treatment on the tumour microenvironment. Both the positive and negative effects of ICI therapy must be explicitly considered to design treatment schedules that are both safe and effective. The combined model and optimal control framework presented in this paper are a first step towards achieving this.

In this work we extended our model of autoimmune myocarditis by connecting it to a tumour growth model. We showed that when the only objective is to delay tumour growth, we recover the standard-of-care treatment schedule. When we consider the effect of ICIs on the tumour and the heart simultaneously, however, we obtain schedules that spread out the doses across the allowed treatment window, striking a balance between keeping tumour growth under control and preventing autoimmune myocarditis from developing.

Our primary aim was to identify a schedule which prevents autoimmune myocarditis and delays tumour growth compared to the treatment-free case. We focused on achieving our objective by finding optimized, rather than optimal, dosing schedules, in the sense that we did not seek to find the global minimum in the payoff landscape. The optimization framework we have presented here can be further developed to not only prevent myocarditis, but to potentially also maximize inhibition of tumour growth by finding the optimal dosing schedule, *i*.*e*. determining the optimal number of doses, dose size and timing of doses. The framework is flexible and thus generally applicable and adaptable to a particular issue of interest related to optimizing cancer treatment.

The models for autoimmune myocarditis and tumour growth used in this work are simple and serve as a starting point from which more complex, biologically detailed models can be built. Including more variables, representing for example specific immune cell types or cytokines to more accurately represent the immunological mechanisms in the model of autoimmune myocarditis, and adding more detail to the tumour compartment, will allow for more realistic simulation of the system. In combination with patient data, the framework developed here for designing dosing schedules could then be used in the clinic to personalize cancer treatment.

Lastly, the heart is not the only organ or tissue impacted by off-target effects of cancer therapy. The gastrointestinal tract, skin and liver are just a few of the other organs that are known to experience side-effects of ICI therapy [30]. To model the range of effects of ICI therapy on the whole body and take a holistic approach to treatment optimization, the heart-tumour model must be extended with compartments representing other parts of the body that are impacted by off-target effects of ICI treatment. Such a modelling framework also allows for the inclusion of other types of therapy, such as radiation therapy, which are administered to patients alone or in combination with ICIs or other therapies. A model which includes all effects of cancer treatment in combination with the optimal control framework we have presented here will be an important step towards the design of personalized dosing schedules for cancer therapies in the clinic.

## Acknowledgments

The authors would like to thank Dr. Liudmila Polonchuk and Dr. Ken Wang for helpful discussion and suggestions.

## Declarations

- Funding: S.A.V. is supported by the EPSRC (EP/L016044/1). REB is supported by a grant from the Simons Foundation (MP-SIP-00001828).
- Conflict of interest/Competing interests: None.
- Data availability: this manuscript has no associated data.
- Authors’ contributions: S.A. van der Vegt undertook the research, and prepared and edited the manuscript. S.L. Waters and R.E. Baker edited the manuscript. S.A. van der Vegt, S. L. Waters and R.E. Baker designed the research.

