## Supplementary Information for "Optimal control of immune checkpoint inhibitor therapy in a heart-tumour model"

#### 1 Optimal control of tumour growth only

We use the simple tumour model defined in Section 2.2 and the optimal control algorithm from Castiglione and Piccoli [9] detailed in Section 3 to find a schedule of ICI doses that delays tumour growth. We construct a payoff function that penalizes tumour growth at any time point in the simulation. The payoff function to be minimized is

$$J = \int_0^{t_f} T_u(t)\mathrm{d}t. \tag{1}$$

In each case, we initialize the model with a growing tumour which is larger than the saddle point but small compared to the carrying capacity, *i.e.* $T_{u,0} = 0.1$ cm$^3$. The convergence threshold, $c_p$, is set to 0.1, and the gradient descent step size, $h$, is equal to 0.01. All model parameters are set to their default values in Tbl. 2.

We set the dose of nivolumab equal to $V_N = 0.25$ grams for monotherapy and combination therapy, and the dose of ipilimumab equal to $V_I = 1.0$ grams for monotherapy and $V_I = 0.1$ grams for combination therapy. We use the same timeline as we did for the two-compartment model, which is shown in Fig. 3.

We initialize the optimization of nivolumab monotherapy, ipilimumab monotherapy and combination therapy with their respective standard-of-care schedules (see Tbl. 1). For all three types of therapy, we find that the optimized treatment schedule is nearly identical to the standard-of-care treatment schedule. For nivolumab monotherapy, the standard-of-care dosing schedule, in days, is $S_0 = [16,\ 30,\ 44,\ 58]$ and the optimized dosing schedule is $S = [16.0,\ 30.0,\ 44.0,\ 58.0]$. For ipilimumab monotherapy, the standard-of-care dosing schedule schedule is $S_0 = [23,\ 44,\ 65]$ and the optimized dosing schedule is $S = [23.0,\ 44.0,\ 65.0]$. For combination therapy, the standard-of-care dosing schedule schedule is $S_{N,0} = [23,\ 44,\ 65,\ 86]$, and $S_{I,0} = [23,\ 44,\ 65]$ and the optimized dosing schedule is $S_N = [23.0,\ 44.0,\ 65.1,\ 86.1]$, and $S_I = [23.0,\ 44.0,\ 65.0]$.

Based on delay of tumour growth alone, the standard-of-care treatment schedule for ICI therapy thus appears to be optimized.

### 2 Exploration of optimal control algorithm parameters

Here we determine designed optimized dosing schedules for nivolumab and ipilimumab monotherapy as well as combination therapy using our heart-tumour model. Here we explore the effect of changing the values of the doses $V_N$ and $V_I$, the length of the treatment window, the step parameter $h$, and the weights $w_1$ and $w_2$ in the payoff function on the obtained optimized dosing schedule. The default value of $V_N$ is $V_N = 0.25$ grams, the default value of $V_I$ is $V_I = 1.0$ grams for ipilimumab monotherapy and $V_I = 0.1$ grams for combination therapy. The final day of the treatment window, $t_{\max}$, is by default day 60 for nivolumab monotherapy, day 70 for ipilimumab monotherapy and day 90 for combination therapy. The default values of the other parameters are $h = 0.01$, and the weights are $w_1 = 1$ cells$^{-1}$, $w_2 = 1$ cm$^{-3}$.

#### 2.1 Nivolumab monotherapy

##### 2.1.1 The effect of changes to $V_N$

We explore the effects of changing the value of $V_N$ on the development of autoimmune myocarditis. We vary the value of $V_N$ and keep all other parameter values at their default values listed in Tbl. 2. We then use the set-up and algorithm as explained in Section 3 to optimize the dosing schedule. We use the standard-of-care schedule as the initial schedule for $V_N = 0.01$ grams, and set the initial schedule for $V_N = m$ grams, $m = 0.02, 0.03, \ldots, 0.27$, to be the optimized schedule for $V_N = m - 0.01$ grams. For $V_N > 0.27$, we set the initial schedule equal $S_0 = [60, 60, 60, 60]$ to speed up convergence.

As shown in Fig. 1, there is a very small range of values of $V_N$ for which autoimmune myocarditis develops when the standard-of-care schedule is applied, but does not develop for an optimized dosing schedule identified by the optimal control algorithm (green). For $V_N \leq 0.24$ grams, the standard-of-care schedule does not cause autoimmune myocarditis. For $V_N \geq 0.28$ grams, the standard-of-care schedule causes autoimmune myocarditis to develop and

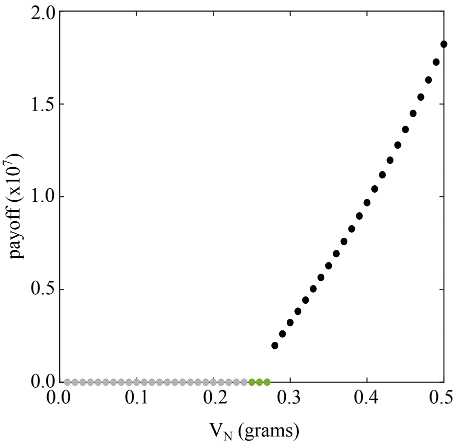


**Fig. 1**: The payoff of the optimized nivolumab dosing schedule as $V_N$ is varied. A grey dot indicates a value of $V_N$ for which autoimmune myocarditis does not develop when a standard-of-care treatment schedule is applied. A green dot indicates a value of $V_N$ for which autoimmune myocarditis does develop when a standard-of-care treatment schedule is applied, but we can find an optimized treatment schedule for which autoimmune myocarditis is prevented. A black dot indicates a value of $V_N$ for which autoimmune myocarditis does develop when a standard-of-care treatment schedule is applied, and this cannot be prevented by optimizing the dosing schedule with the parameter values we consider (see Tbl. 2).

a myocarditis-free solution cannot be identified. That leaves the range $0.25 \leq V_N \leq 0.27$ grams for which we can optimize the dosing schedule such that it does not cause autoimmune myocarditis where the standard-of-care dosing schedule does, within the given constraints, further motivating our choice for $V_N = 0.25$ grams in the main text.

##### 2.1.2 The effect of changes to the final dosing day, $t_{\max}$

We now explore the effects of increasing the length of the treatment window on the dosing schedule. We set all other parameters to the values in Tbl. 2 and let the final day of the dosing window, $t_{\max}$, vary from day 60 to day 110. We use the standard-of-care schedule as the initial schedule for $t_{\max} = 60$ days, and thereafter set the initial schedule for $t_{\max} = m$ days, $m = 61, 62, \ldots, 110$, to be the optimized schedule for $t_{\max} = m - 1$ days.

Fig. 2 shows the results. Note that for every value of $t_{\max}$ the last dose in the schedule consists of two doses administered simultaneously. As the length of the treatment window is extended to $t_{\max} = 82$ days, the last doses of the schedule are delayed to the last day of the dosing window. For treatment windows where $t_{\max} > 82$ days, the last two doses are consistently administered on day 82. This can be explained by the fact that dosing later than day 82

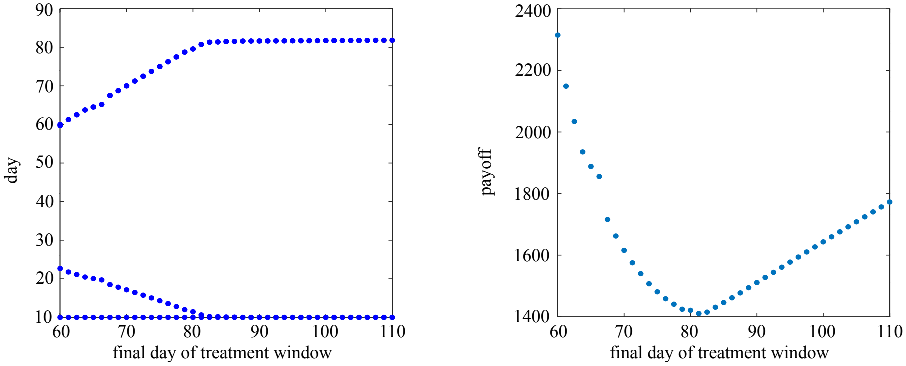


**Fig. 2**: Optimized nivolumab dosing schedule (left) and payoff (right) for different lengths of the treatment window. Parameter values in Tbl. 2 are used unless otherwise noted.

would leave too much time between doses during which the tumour can grow uninhibited. The first two doses in the optimized part of the dosing schedule are administered separately early in the treatment window for shorter treatment windows, and simultaneously on day 10 from $t_{\max} = 82$ days. Overall, we see that the trend of dosing early as well as late in the treatment window such that tumour growth is inhibited by regular dosing and sufficient time is left between doses to prevent autoimmune myocarditis holds as the treatment window is extended.

##### 2.1.3 The effect of changes to $h$

We now vary the step size parameter $h$ to ensure robustness of the results with respect to this algorithm parameter. We keep all other parameters at the default values listed in Tbl. 2 and initialize with the standard-of-care dosing schedule.

The results in Tbl. 1 show for all values of $h$ used here, we see that the optimized dosing schedule for nivolumab monotherapy consists of two early doses administered before day 30 and two late doses administered on day 47 or thereafter. None of the dosing schedules in Tbl. 1 cause autoimmune myocarditis to develop and all inhibit tumour growth, so although the exact timing of doses varies somewhat, the main objectives of inhibiting tumour growth and preventing autoimmune myocarditis are achieved for all values of $h$.

##### 2.1.4 The effect of payoff function weights

Lastly, we explore the effects of varying the weights in the payoff function. The payoff function used for all types of ICI therapy is

| $h$ | Schedule nivolumab | Payoff |
|---|---|---|
| 0.001 | 10.0, 28.4, 47.9, 60.0 | $3.9415 \times 10^3$ |
| 0.005 | 10.0, 22.5, 60.0, 60.0 | $2.2870 \times 10^3$ |
| 0.01 | 10.0, 22.7, 59.7, 60.0 | $2.3148 \times 10^3$ |
| 0.05 | 10.0, 10.0, 60.0, 60.0 | $1.8514 \times 10^3$ |
| 0.1 | 10.0, 10.0, 60.0, 60.0 | $1.8514 \times 10^3$ |

**Table 1**: The optimized schedule of nivolumab doses and the payoff vary as the size of the step size parameter, $h$, is varied. None of the dosing schedules in this table cause autoimmune myocarditis to develop.

| $w_2$ | Schedule nivolumab | Payoff |
|---|---|---|
| 0.001 | 10.0, 22.6, 59.7, 60.0 | $2.2770 \times 10^3$ |
| 0.1 | 10.0, 22.6, 59.7, 60.0 | $2.2808 \times 10^3$ |
| 1 | 10.0, 22.7, 59.7, 60.0 | $2.3148 \times 10^3$ |
| 10 | 10.0, 23.1, 60.0, 60.0 | $2.6256 \times 10^3$ |
| 100 | 10.0, 26.6, 60.0, 60.0 | $5.8737 \times 10^3$ |

**Table 2**: The optimized schedule of nivolumab doses and the payoff vary as the weight $w_2$ is varied. None of the dosing schedules in this table cause autoimmune myocarditis to develop.

$$J = \int_0^{t_f} [w_1 C(t) + w_2 T_u(t)] \, \mathrm{d}t. \tag{2}$$

Since only the relative values of the weights matters, it suffices to vary only $w_2$ and keep $w_1 = 1$ cells$^{-1}$. We vary $w_2$ across the range [0.01, 0.1, 1, 10, 100] cm$^{-3}$ and optimize the treatment schedule for each value in this range. We keep all other parameters at the values listed in Tbl. 2 and initialize the dosing schedule as the standard-of-care schedule.

Tbl. 2 shows that for each value of $w_2$ the optimized dosing schedule is qualitatively the same. The payoff also remains on the same order of magnitude. Because the weights appear in the payoff function, we would expect the payoff to vary somewhat as different parts of the payoff function are prioritized and weighted differently. For all values of $w_2$, the algorithm can identify a dosing schedule which does not cause autoimmune myocarditis to develop and inhibits tumour growth, although the optimized schedule differ somewhat for different values of $w_2$. We can thus conclude that the optimization of nivolumab dosing schedules is robust to changes in the weights in the payoff function.

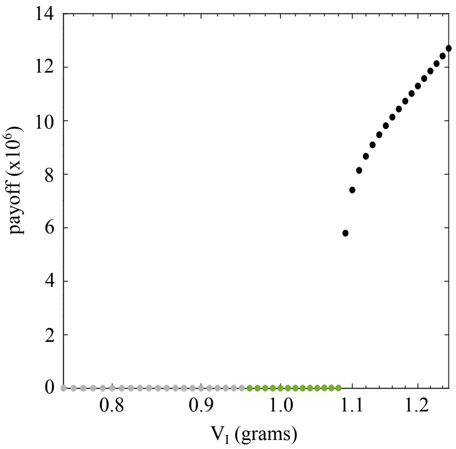


**Fig. 3**: The payoff of the optimized ipilimumab dosing schedule as $V_I$ is varied. A grey dot indicates a value of $V_I$ for which autoimmune myocarditis does not develop when a standard-of-care treatment schedule is applied. A green dot indicates a value of $V_I$ for which autoimmune myocarditis does develop when a standard-of-care treatment schedule is applied, but we can find an optimized treatment schedule for which autoimmune myocarditis is prevented. A black dot indicates a value of $V_I$ for which autoimmune myocarditis does develop when a standard-of-care treatment schedule is applied, and this cannot be prevented by optimizing the dosing schedule.

#### 2.2 Ipilimumab monotherapy

##### 2.2.1 The effect of changes to $V_I$

We vary the value of $V_I$ to establish the range of values for which autoimmune myocarditis develops as a side-effect of the standard-of-care schedule, but can be prevented by an optimized dosing schedule. We set all other parameters to the values listed in Tbl. 2. We use the standard-of-care schedule as the initial schedule for $V_I = 0.75$ grams, and thereafter set the initial schedule for $V_I = m$ grams, $m = 0.76, 0.77, \ldots, 1.25$, to be the optimized schedule for $V_I = m - 0.01$ grams.

Fig. 3 shows that we again obtain three regions with different outcomes. For $V_I \leq 0.95$ grams, autoimmune myocarditis does not develop when the standard-of-care schedule is applied. For $V_I \geq 1.09$ grams, autoimmune myocarditis is a side-effect when the standard-of-care schedule is applied, and it cannot be prevented by an optimized schedule. This leaves the range $0.96 \leq V_I \leq 1.08$ grams for which autoimmune myocarditis is a side-effect of the standard-of-care treatment schedule, but it can be prevented by an optimized dosing schedule.

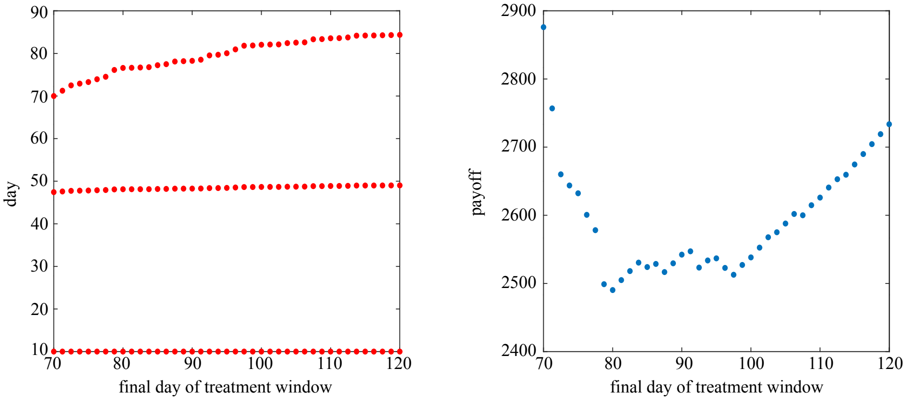


**Fig. 4**: Optimized ipilimumab dosing schedule (left) and payoff (right) for different lengths of the treatment window. Parameter values listed in Tbl. 2 are used unless otherwise noted.

##### 2.2.2 The effect of changes to the final dosing day

We now turn to the effect of the length of the treatment window on the optimization of treatment schedules for ipilimumab monotherapy. The last day of the treatment window, $t_{\max}$, is day 70 by default. We now let $t_{\max}$ increase from 70 days to 120 days, while keeping all parameter values as listed in Tbl. 2. We use the standard-of-care schedule as the initial schedule for $t_{\max} = 70$ days, and thereafter set the initial schedule for $t_{\max} = m$ days, $m = 71, 72, \ldots, 120$, to be the optimized schedule for $t_{\max} = m - 1$ days.

Fig. 4 shows that the results of the optimization are qualitatively robust to changes in the length of the dosing window. The second and third doses, counting the dose administered on day 2 as the first dose, are administered approximately on the same day, regardless of the length of the treatment window. The fourth dose ipilimumab is administered later as the treatment window increases. This even spread of the second, third and fourth doses over the treatment window controls tumour growth while leaving enough time between doses for immune cell counts to decrease, thus preventing autoimmune myocarditis.

##### 2.2.3 The effect of $h$

We explore the robustness of the optimization results for ipilimumab monotherapy with respect to changes in the step parameter $h$. We keep all other parameters at the values listed in Tbl. 2 while varying $h$ from 0.001 to 0.1. We initialize the dosing schedule at the standard-of-care schedule.

The results in Tbl. 3 show that the value of $h$ does not qualitatively affect the results of the optimization. For all values of $h$ used here, we see that the optimized dosing schedule for ipilimumab monotherapy consists of one early dose, one dose around day 47 and one late dose on day 70, thus covering

| $h$ | Schedule ipilimumab | Payoff |
|---|---|---|
| 0.001 | 15.1, 46.6, 70.0 | $3.8528 \times 10^3$ |
| 0.005 | 13.7, 46.7, 70.0 | $3.4658 \times 10^3$ |
| 0.01 | 10.0, 47.4, 70.0 | $2.8761 \times 10^3$ |
| 0.05 | 10.0, 47.4, 70.0 | $2.8759 \times 10^3$ |
| 0.1 | 10.0, 47.5, 70.0 | $2.8749 \times 10^3$ |

**Table 3**: The schedule of ipilimumab doses and the payoff vary as the size of the step size parameter, $h$, is varied. None of the dosing schedules in this table cause autoimmune myocarditis to develop.

the whole treatment window. None of the dosing schedules in Tbl. 3 cause autoimmune myocarditis to develop and all inhibit tumour growth.

##### 2.2.4 The effect of payoff function weights

Lastly, we explore the effects of varying the weights in the payoff function on the optimization of ipilimumab monotherapy. The payoff function used for all types of therapy is

$$J = \int_0^{t_f} [w_1 C(t) + w_2 T_u(t)] \,\mathrm{d}t. \tag{3}$$

Since only the relative values of the weights matters, it suffices to vary only $w_2$, while keeping $w_1 = 1$ cells$^{-1}$. We vary $w_2$ across the range $[0.01,\ 0.1,\ 1,\ 10,\ 100]$ cm$^{-3}$ and optimize the treatment schedule for each value in this range. We keep all other parameters at the values listed in Tbl. 2 and initialize the dosing schedule as the standard-of-care schedule.

Tbl. 4 shows that for each value of $w_2$, the optimized dosing schedules remain qualitatively similar. Because the weights appear in the payoff function, we would expect the payoff to vary somewhat as different parts of the payoff function are prioritized and weighted differently, and this is reflected in Tbl. 4. For all values of $w_2$ the algorithm identifies a dosing schedule which does not cause autoimmune myocarditis to develop and inhibits tumour growth. We can thus conclude that the optimization of ipilimumab dosing schedules is robust to changes in the weights in the payoff function.

#### 2.3 Combination therapy

##### 2.3.1 The effect of changes to $V_N$ and $V_I$

We explore the effects of varying the values of $V_N$ and $V_I$ simultaneously over the ranges 0.1-0.3 grams and 0.0-0.3 grams, respectively. We keep all other parameters at the values in Tbl. 2. To speed up the optimization, for each value of $V_N$ we initialize the dosing schedule at the standard-of-care schedule of $V_I = 0$ grams, and thereafter set the initial schedule for $V_I = m$ grams, $m = 0.01, 0.02, \ldots, 0.30$, to be the optimized schedule for $V_I = m - 0.01$ grams.

| $w_2$ | Schedule ipilimumab | Payoff |
|---|---|---|
| 0.01 | 10.0, 47.4, 70.0 | $0.2353 \times 10^4$ |
| 0.1 | 10.0, 47.4, 70.0 | $0.2400 \times 10^4$ |
| 1 | 10.0, 47.4, 70.0 | $0.2876 \times 10^4$ |
| 10 | 10.0, 47.5, 70.0 | $0.7590 \times 10^4$ |
| 100 | 16.2, 46.6, 70.0 | $5.5136 \times 10^4$ |

**Table 4**: The schedule of ipilimumab doses and the payoff vary as the weight $w_2$ is varied. None of the dosing schedules in this table cause autoimmune myocarditis to develop.

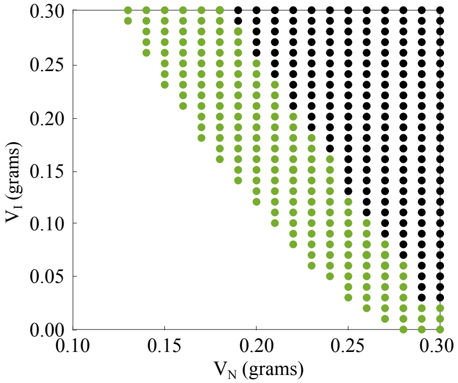


**Fig. 5**: Treatment outcome of the optimized combination therapy dosing schedule as $V_N$ and $V_I$ are varied simultaneously. In the white region of values of $V_N$ and $V_I$ autoimmune myocarditis never develops. In the green region of values of $V_N$ and $V_I$ autoimmune myocarditis is a possible side-effect of combination therapy but it can be prevented by an optimized dosing schedule. In the black region of values of $V_N$ and $V_I$ autoimmune myocarditis cannot be prevented as a side-effect.

We find again that there are three types of outcome as we vary $V_N$ and $V_I$. The white area in Fig. 5 indicates the values for which autoimmune myocarditis does not develop as a side-effect of the standard-of-care treatment schedule. The green area indicates the values for which myocarditis does develop as side-effect of the standard-of-care treatment schedule, but we can optimize the treatment schedule such that we prevent this. The black area indicates those values for which autoimmune myocarditis cannot be prevented within the constraints set here.

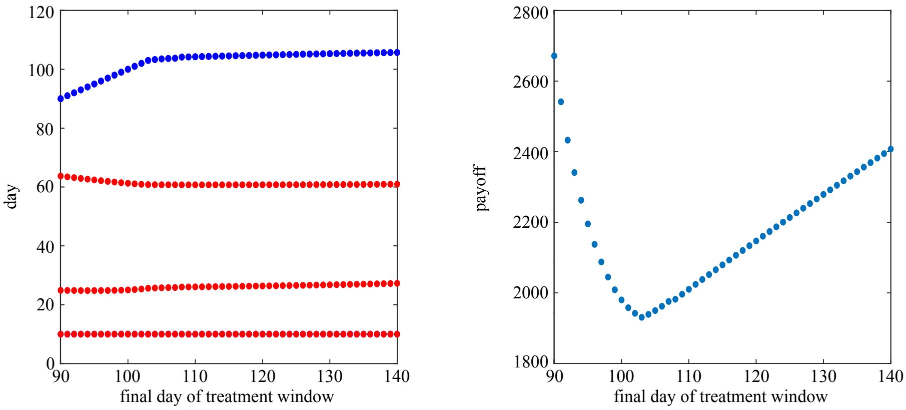


**Fig. 6**: Optimized combination therapy dosing schedule (left) and payoff (right) for different lengths of the treatment window. Parameter values listed in Tbl. 2 are used unless otherwise noted. For the optimization of the dosing schedule for each length of treatment window, the initial dosing schedule, $S_0$, is set to be the standard-of-care dosing schedule.

##### 2.3.2 The effect of extending the treatment window

We now consider the effect that extending the treatment window has on the optimization of combination therapy treatment schedules. All parameters are set to the values listed in Tbl. 2. The final day of the treatment window, $t_{\max}$, is varied from day 90 to 140. To speed up the optimization, we use the standard-of-care schedule as the initial schedule for $t_{\max} = 90$ days, and thereafter set the initial schedule for $t_{\max} = m$ days, $m = 91, 92, \ldots, 140$, to be the optimized schedule for $t_{\max} = m - 1$ days.

Fig. 6 shows the results. Note that the last dose of nivolumab (blue) consists of three doses administered simultaneously, and the second dose, administered on day 10 combines one dose of nivolumab and one dose of ipilimumab. As Fig. 6 shows, the optimization of the treatment schedule for combination therapy is robust to changes in the length of the treatment window. As the treatment window is extended, the optimized schedule does not change qualitatively. The final three doses of nivolumab, which are administered simultaneously, are delayed slightly but the timing of the other doses remains mostly constant as $t_{\max}$ is varied. The payoff of each of these schedules also does not change significantly taking into account that a longer simulation is expected to lead to a higher payoff as the integral is computed over a longer time span.

##### 2.3.3 The effect of $h$

Next we investigate robustness of the optimization results to varying the step size parameter $h$. To speed up the optimization of combination therapy dosing schedules, we set $h = 0.1$ initially and then change to $h = 0.01$ once the payoff is less than $10^4$. Here we vary the value of $h$ once the payoff is less than $10^4$ and

| $h$ | Schedule nivolumab | Schedule ipilimumab | Payoff |
|---|---|---|---|
| 0.001 | 10.0, 77.2, 90.0, 90.0 | 10.0, 25.9, 69.1 | 5647.9 |
| 0.005 | 10.0, 89.8, 90.0, 90.0 | 10.0, 24.9, 63.8 | 2695.7 |
| 0.01 | 10.0, 90.0, 90.0, 90.0 | 10.0, 24.9, 63.7 | 2671.5 |
| 0.05 | 10.0, 90.0, 90.0, 90.0 | 10.0, 24.25, 58.3 | 2558.6 |
| 0.1 | 10.0, 90.0, 90.0, 90.0 | 10.0, 23.4, 52.9 | 2480.4 |

**Table 5**: The schedule of combination therapy and the payoff vary as the size of the step size parameter, $h$, is varied. None of the dosing schedules in this table cause autoimmune myocarditis to develop.

leave $h = 0.1$ above this threshold. All other parameters are set to the values listed in Tbl. 2 and we initialize the dosing schedule as the standard-of-care schedule for all $h$.

None of the schedules in Tbl. 5 cause autoimmune myocarditis to develop. Furthermore, the schedules do not change qualitatively as the value of $h$ is varied. The results of the optimization of combination therapy are thus robust to our choice of $h$.

##### 2.3.4 The effect of payoff function weights

Lastly, we explore the effect of changing the weights in the payoff function on the optimization results. The payoff function used for all types of therapy is

$$J = \int_0^{t_f} [w_1 C(t) + w_2 T_u(t)] \, \mathrm{d}t. \tag{4}$$

Since only the relative values of the weights matters, it suffices to vary only $w_2$ and keep $w_1 = 1$ cells$^{-1}$. We vary $w_2$ across the range $[0.01,\ 0.1,\ 1,\ 10,\ 100]$ cm$^{-3}$ and optimize the treatment schedule for each value of $w_2$. We keep all other parameters at the values listed in Tbl. 2 and initialize the dosing schedule as the standard-of-care schedule.

Results in Tbl. 6 show that we need to take some care when picking the weights in the payoff function for combination therapy. For $w_2 = 0.01$ the optimized schedule still causes autoimmune myocarditis to develop. The evolution of the dosing schedule and the payoff over the iterations of the gradient descent algorithm with this value of $w_2$ show that, up to convergence, the evolution of the dosing schedule is qualitatively the same as happens in the cases where the algorithm does find a myocarditis-free solution, e.g. with the default values $w_1 = 1$ cells$^{-1}$ and $w_2 = 1$ cells$^{-1}$. The problem thus appears to be that the threshold for convergence is met far away from a local minimum in the payoff landscape, likely due to the flatness of the payoff landscape in a large neighborhood around the minimum. This indicates that careful calibration of the step parameter $h$, the payoff functions weights $w_1$ and $w_2$, and the convergence threshold $c_p$ is necessary to ensure convergence to a myocarditis-free solution.

| $w_2$ | Schedule nivolumab | Schedule ipilimumab | Payoff |
|---|---|---|---|
| 0.01 | 10.0, 68.8, 90.0, 90.0 | 14.5, 32.2, 71.6 | $1.8297 \times 10^5$ |
| 0.1 | 10.0, 90.0, 90.0, 90.0 | 10.0, 24.9, 63.7 | $2.6407 \times 10^3$ |
| 1 | 10.0, 90.0, 90.0, 90.0 | 10.0, 24.9, 63.7 | $2.6716 \times 10^3$ |
| 10 | 10.0, 89.8, 90.0, 90.0 | 10.0, 25.0, 63.7 | $3.0039 \times 10^3$ |
| 100 | 10.0, 80.4, 90.0, 90.0 | 10.0, 25.7, 67.2 | $7.4791 \times 10^3$ |

**Table 6**: The schedule of combination therapy and the payoff vary as the weight $w_2$ is varied. The optimized schedule for $w_2 = 0.01$ is the only schedule which causes autoimmune myocarditis to develop.

In this appendix we have shown that for nivolumab and ipilimumab monotherapy, the results of optimization of the dosing schedule are robust to changes in the length of the treatment window, the step parameter $h$, and the weights in the payoff function $w_1$ and $w_2$. For combination therapy, the optimized schedule is not significantly affected by changes to length of the treatment window or the step parameter $h$ if all other parameters are set to their default values, but care should be taken when setting the values of $h$ in combination with certain payoff function weights. The weights of the payoff function are ideally set by the biological context of the optimization problem and we have shown that for certain values of the weights the algorithm fails to converge to a myocarditis-free solution even though a myocarditis-free solution is possible within the set constraints. This is due to the algorithm hitting the convergence threshold before it reaches the myocarditis-free solution. This suggests that the default value of $h$ that we have used throughout this work may not be appropriate for all values of the weights and should thus be carefully considered.
